# The impact of stroma on the discovery of molecular subtypes and prognostic gene signatures in serous ovarian cancer

**DOI:** 10.1101/496406

**Authors:** Matthew Schwede, Levi Waldron, Samuel C. Mok, Wei Wei, Azfar Basunia, Melissa A. Merritt, Giovanni Parmigiani, David Harrington, John Quackenbush, Michael J. Birrer, Aedín C. Culhane

## Abstract

**Purpose:** Recent efforts to improve outcomes for high-grade serous ovarian cancer, a leading cause of cancer death in women, have focused on identifying molecular subtypes and prognostic gene signatures, but existing subtypes have poor cross-study robustness. We tested the contribution of cell admixture in published ovarian cancer molecular subtypes and prognostic gene signatures.

**Experimental Design:** Public gene expression data, two molecular subtype classifications, and 61 published gene signatures of ovarian cancer were examined. Using microdissected data, we developed gene signatures of ovarian tumor and stroma. Computational simulations of increasing stromal cell proportion were performed by mixing gene expression profiles of paired microdissected ovarian tumor and stroma.

**Results:** Established ovarian cancer molecular subtypes are strongly associated with the cell admixture. Tumors were classified as different molecular subtypes in simulations, when the percentage of stromal cells increased. Stromal gene expression in bulk tumor was weakly prognostic, and in one dataset, increased stroma was associated with anatomic sampling location. Five published prognostic gene signatures were no longer prognostic in a multivariate model that adjusted for stromal content alone.

**Conclusions:** The discovery that molecular subtypes of high grade serous ovarian cancer is influenced by cell admixture, and stromal cell gene expression is crucial for interpretation and reproduction of ovarian cancer molecular subtypes and gene signatures derived from bulk tissue. Single cell analysis may be required to refine the molecular subtypes of high grade serous ovarian cancer. Because stroma proportion was weakly prognostic, elucidating the role of the tumor microenvironment’s components will be important.

**Translational relevance:** Ovarian cancer is a leading cause of cancer death in women in the United States. Although the tumor responds to standard therapy for the majority of patients, it frequently recurs and becomes drug-resistant. Recent efforts have focused on identifying molecular subtypes and prognostic gene signatures of ovarian cancer in order to tailor therapy and improve outcomes. This study demonstrates that molecular subtype identification depends on the ratio of tumor to stroma within the specimen. We show that the specific anatomic location of the biopsy may influence the proportion of stromal involvement and potentially the resulting gene expression pattern. It will be crucial for these factors to be taken into consideration when interpreting and reproducing ovarian cancer molecular subtypes and gene signatures derived using bulk tissue and single cells. Furthermore, it will be important to define the relative proportions of stromal cells and model their prognostic importance in the tumor microenvironment.

## Introduction

There have been several attempts to identify similar molecular subtypes and prognostic genomic signatures for advanced-stage serous ovarian cancers (1–4). The first large study to report molecular subtypes studied mostly serous tumors and identified a subtype that was associated with poor overall survival, C1 (1). Subsequently, The Cancer Genome Atlas (TCGA) reported four similar subtypes in high grade serous ovarian cancer (HGSOC) but found no difference in clinical outcome between the subtypes (2), which led researchers to question the clinical utility of these classifications (5). Subtype predictions using the TCGA classification are not consistently robust, sometimes producing overlapping clusters, with most tumors assigned to more than one subtype (6,7). A recent comprehensive meta-analysis could not robustly classify subtypes across datasets (8) and led to alternative subtype classifications that were associated with survival, tumor purity, age, and lymphocyte infiltration (8).

Molecular subtype and biomarker discovery in HGSOC has mostly been performed on gene expression profiles of bulk tumors that may include fibroblasts, immune cells, and endothelial cells (9) and has not accounted for tumor purity, which can be highly variable within and between studies. Stromal content is variable in two of the largest gene expression profiling studies (1,2). In the Australian Ovarian Cancer Study (AOCS), 40% of tumors in the molecular subtype C1 had low tumor percentage (≤ 50%) compared with 9% in the other molecular subtypes combined (1). In the TCGA study, the proportion of stromal cells exceeded 50% in about one percent of tumors, and the median proportion of stromal cells was 10% (2). Cells in the tumor microenvironment are important for chemo-sensitivity (10), and genes expressed by cells in the tumor microenvironment are prognostic in ovarian cancer (4,11–13). For example, activation of Smad signaling in cancer associated fibroblasts in the tumor microenvironment is associated poor patient survival (13). The the ratio of tumor to stroma may also be an independent prognostic indicator in several cancers, including breast (14), colon (15), and ovarian cancer (16).

However, a bulk molecular profile is the sum of the gene counts in an admixture of carcinoma and stromal cells, which may confound inference of properties of the malignant epithelial cell itself. Single-cell gene expression profiling may resolve many of these issues, but the extent to which non-malignant cell admixture has influenced the field to date is unknown.

We systematically explored the extent to which stromal cell admixture in tumor samples influences molecular subtype and prognostic gene signature discovery (Figure 1). We hypothesized that if unadjusted in statistical analyses, variance in percent stromal cells may lead to false discovery of tumor molecular subtypes, low precision of tumor classification, and reduced power to detect true molecular subtypes and prognostic signatures. We tested this by identifying genes that were differentially expressed between microdissected tumor and stromal cells and examining whether these genes were enriched in published serous ovarian cancer subtypes and prognostic gene expression signatures. We show that percent stromal content itself is weakly prognostic and investigate whether published prognostic ovarian cancer gene signatures simply capture the prognostic effect of the ratio of tumor to stroma or extract differential cancer cell gene expression.

**Figure 1:**
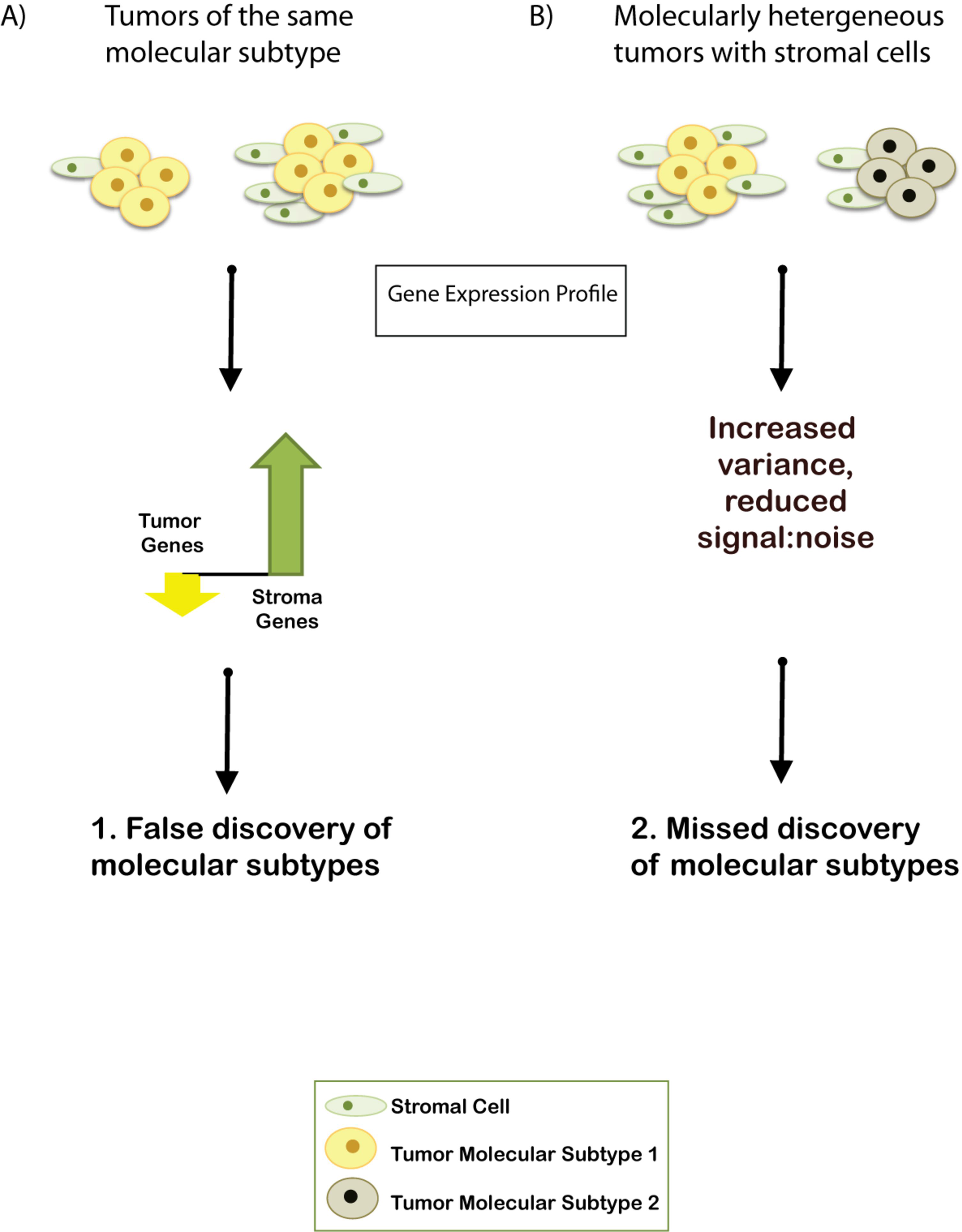
Variability in stromal proportions may present challenges in gene expression analysis. A) If two tumors of the same molecular subtype were sampled with different proportions of stromal cells, one would observe an increase in stroma-associated gene expression with a concurrent decrease in tumor-associated gene expression. This sampling issue may result in false discovery of molecular subtypes due to variable gene expression from epithelial and stromal genes. B) Different proportions of stromal cells will increase noise in analysis, which may cause researchers not to discover underlying molecular subtypes.

## Materials and Methods

### Data

We downloaded two datasets from the Australian Ovarian Cancer Study (AOCS) (2): 1) gene expression profiles of high-grade (G2, G3) ovarian adenocarcinomas (n=215; 11 endometrioid, 203 serous, one undifferentiated) that were previously classified as high grade molecular subtypes C1, C2, C4, or C5 (the “AOCS dataset”, GEO GSE9891) and 2) a subset of four tumors that had matched microdissected tumor and stroma (the “AOCS microdissected dataset”, n=8, GEO GSE9890). In the AOCS microdissected dataset, only tumors of the C1/stromal subtype were selected for analysis (1). In the AOCS dataset, we excluded low malignant potential tumors (C3 and C6 subtypes, n=36) and unclassified tumors (n=4). The AOCS and AOCS microdissected datasets were normalized using frozen robust multi-array analysis (fRMA) (17). The validation ovarian cancer gene expression datasets included: 1) Affymetrix gene expression profiles of pairs (n=38) of microdissected tumor and stroma of HGSOC from women hospitalized at the Massachusetts General Hospital (GSE40595, the “MGH dataset”) (18,19), 2) the Cancer Genome Atlas (TCGA) ovarian cancer dataset (n=518, downloaded November 24, 2010) in which all but one tumor was high-grade serous (2), and 3) the aggregated datasets in the curatedOvarianData Bioconductor package (20). Datasets in the curatedOvarianData package are listed in Table S1. Non-ovarian validation data included microarray datasets of breast and prostate cancer that contained microdissected stroma and tumor or reported the percent stromal content of bulk tumor. These included microdissected paired tumor epithelium and stroma (n=34 pairs) from non-inflammatory breast cancer (GSE5847, Boersma et al.) (21); breast cancer epithelium and stroma pairs (n=28 pairs) from a study of invasive breast cancer (GSE10797, Casey et al.) (22); and two prostate cancer datasets with known percent stromal content (GSE17951, n=136; GSE8218, n=109) (9). All validation datasets were downloaded from GEO, except that the MGH dataset was received directly from the authors, and normalized using RMA (23,24).

### Statistical analysis

Analyses were performed using R and Bioconductor, versions 2.15 and 2.10, respectively, or later.

### Gene signatures of tumor and stroma

Genome-wide differential gene expression analyses were performed using a moderated *t*-test, and p-values were adjusted for multiple testing using the Benjamini-Hochberg method (25) using the Bioconductor package limma (26). For paired microdissected stroma and tumor samples, a paired moderated *t*-test, with a p-value cutoff of 0.01 was used. A separate analysis of the AOCS datasets with a less stringent cutoff (p<0.05) was performed to identify a broader set of tumor and stromal genes. Genes that were differentially expressed between each AOCS or TCGA molecular subtype pair were detected using an unpaired test. In differential gene expression analysis of the MGH dataset, a more stringent p-value cut-off was applied to generate gene lists of similar length. Hierarchical clustering was performed using Pearson correlations with average linkage clustering (27). We also used these clustering methods when generating heatmaps using scores of immune cell signatures that have been previously described (28). Using binary on/off calls (29) of gene expression in the MGH dataset, we generated tissue-specific marker genes using sensitivity and specificity. These tissue-specific genes were defined using an area under the ROC curve (30) of ≥0.9 for 1) tumor (n=38) 2) paired “tumor stroma” (stroma that is adjacent to tumor, n=38) or 3) unmatched normal ovary stroma (n=10). To generate a single gene signature covariate in statistical models, a gene signature score was computed using a weighted sum approach as previously described (31). Positive weights (+1) were applied to genes up-regulated in poor prognosis tumors, BRCA mutation-like tumors, metastases, an angiogenic subtype, or malignant tumors. Negative weights (−1) were applied to the other genes in the signature. Linear regression was used to test for association between continuous variables and percent stromal content. Fisher’s exact test was used to test for overlap between gene sets, with the background number of genes equal to the number of genes on the platform (for example the AOCS n=18,769).

### Molecular subtype prediction

The gene lists reported to predict AOCS and TCGA molecular subtypes were downloaded as online publication supplements (1,2), curated, and submitted to GeneSigDB (32). The gene lists were used to build a subtype classifier using the single sample predictor (SSP) approach (33). The centroids of gene expression profiles of these genes in each of the four molecular subtypes in the AOCS study (C1, C2, C4, C5) (1) and TCGA study (mesenchymal, immunoreactive, differentiated, proliferative) (2) were calculated (Figure S2). Cases (tumor or stroma) were assigned a molecular subtype based on the highest Pearson correlation coefficients to each subtype’s centroid. Cases were not classified to a molecular subtype (“unclassified”) if its correlation to all subtypes was less than 0.7.

### Computational mixing of tumor and stroma fractions

Paired microdissected tumor and stroma (n=38) were computationally mixed in increasing increments of 10%, as described previously (34). The expression value of each tumor gene (G_t_) was mixed with gene expression value of its adjacent stroma (G_s_) using the linear combinations *m*G_t_ + (1-*m*)G_s_. Mixing parameter *m* is the proportion of tumor and 1-*m* is the proportion of stroma, incremented by 0.1 from 0 to 1. This was performed for all genes in the molecular subtype classifier to produce a computationally simulated mixture of tumor and stroma. The assumption of linearity has been previously supported by comparing the computational values to those observed in gene expression of co-cultured cells for the breast molecular subtype PAM50 gene signature (34).

### Survival analysis

Cox proportional hazards regression was used to test for association between continuous variables and survival variables (relapse-free survival (RFS), overall survival (OS)). In survival analyses of gene signatures in independent datasets, p-values were not corrected for multiple testing because we wished to replicate the expected results as analyzed in their original studies. The prognostic power of the tumor and stroma gene signatures were assessed for concordance of risk scores with overall patient survival using Uno’s version of the Concordance Index (or C-Index). The C-Index represents the probability that a patient predicted to be at lower risk than another patient will survive longer; its expected value is 0.5 for random predictions and 1 for a perfect risk model.

### Exploratory data analysis of stromal genes and TCGA Immune Landscape Signatures

Clinical data (overall survival, debulking, etc.), stromal fraction, leukocyte fraction, the five core immune signatures (wound healing, macrophage regulation, lymphocyte infiltration, interferon gamma response, and TGF-β response), CIBERSORT estimates of >25 immune cells, and mutation load in TCGA HGSOC tumors are available at https://isb-cgc.shinyapps.io/shiny-iatlas/, on the ISB cloud. These are well described as part of the TCGA immune landscape paper (28). Genes with a positive or negative Spearman correlation greater than 0.7 to covariates were visualized using a heatmap to explore associations.

## Results

We investigated the contribution of non-tumor cells’ gene expression to HGSOC molecular subtypes using three different approaches, and we found that the expression of stroma-associated genes substantially affects subtype classification.

### Ovarian cancer molecular subtype gene signatures are enriched in genes expressed in stroma

We investigated the contribution of non-epithelial cell gene expression to HGSOC molecular subtypes by comparing their gene signatures to those of microdissected tumor and stroma. We developed gene signatures of high-grade serous ovarian epithelial carcinoma and adjacent tumor stroma from paired laser microdissected tissue from two independent studies. A 688-gene signature discriminated tumor and stroma paired tissue (n=8) microdissected from tumors of the C1/stromal subtype in the AOCS study (p<0.01, paired t-test with modified variance (26) and FDR correction (25); Table S2, Figure 2). This AOCS tumor-stroma gene signature contained 461 and 227 unique genes with increased expression in C1 stroma and tumor, which are referred to as the “stromal gene set” and “tumor gene set”, respectively. A list with a larger number of genes extracted using lower differential expression stringency (p<0.05) is available in Table S2. In an independent analysis of paired microdissected tumor and stroma dissected from tumors of 38 patients in a Massachusetts General Hospital (MGH) study (18), we identified a second gene signature that discriminated tumor and stroma of high grade serous ovarian cancer. This MGH tumor-stroma gene signature of 519 unique genes contained 429 and 90 genes that had higher expression in stroma and tumor, respectively (Figure S1). Despite the small sample size from which the AOCS stromal signature was derived, the MGH signature had significant overlap with it; 23/81 genes over-expressed in tumor (Fisher’s exact test p<10^−24^) and 125/431 genes over-expressed in stroma (p<10^−99^). The concordance of these data was supported by the observation that genes in the AOCS signature correctly discriminated all but one of the microdissected tumor and stroma MGH specimens using unsupervised hierarchical cluster analysis (Figure S1).

**Figure 2:**
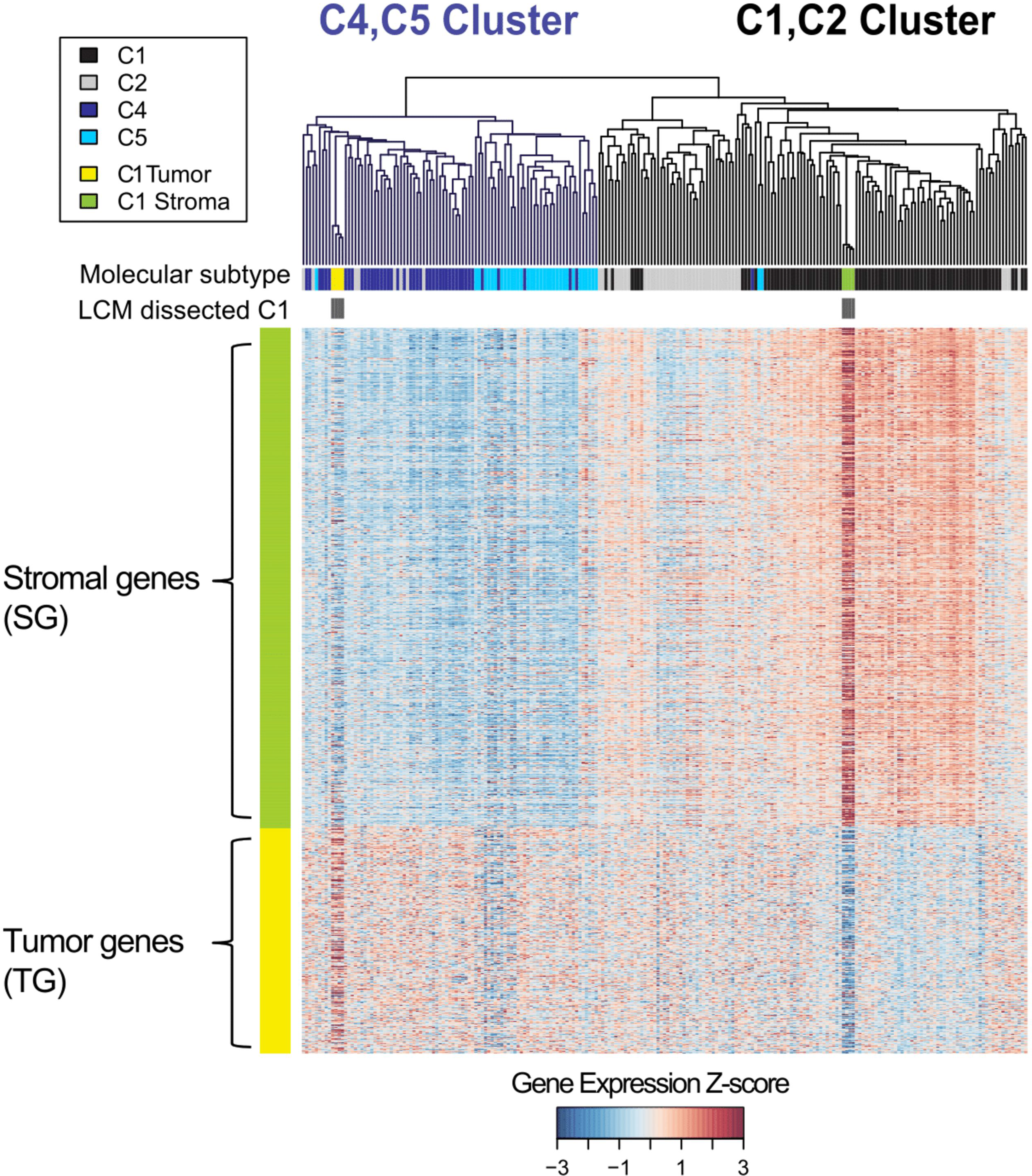
Aocs molecular subtypes can be distinguished by stromal and tumor gene expression. Analysis of gene expression profiles of microdissected epithelial tumor cells and stroma from four C1 tumors identified a 688- gene ovarian tumor-stroma gene signature (FDR adjusted p<0.01). The signature contained 461 genes that were over-expressed in microdissected stroma and 227 genes with increased expression in epithelial tumor cells, and these distinguished ovarian tumor and stroma in an independent MGH dataset (Fig S1). The heatmap shows the tumor and stromal gene expression profiles in the AOCS’s HGSOC tumors and microdissected C1 stroma and tumor samples. C1 microdissected stroma samples clustered with C1 bulk tumors, while the C1 microdissected tumor samples clustered with C4 bulk tumors.

Using these gene signatures, we first investigated the contribution of stromal cell gene expression to the HGSOC molecular subtype using unsupervised cluster analysis. The 688-gene AOCS tumor-stroma gene signature partitioned the AOCS high-grade serous tumors (n=215) into two clusters (Figure 2): those with low and high stromal gene expression. These two clusters could be cut to form four clusters, which largely reflected the four AOCS HGSOC molecular subtypes: C4/CA125, C5/MYCN deregulation (35), C2/immunoreactive, and C1/stromal. We observed that matched microdissected tumor and stroma samples did not cluster with its matched bulk tumor. Instead microdissected C1 tumors clustered with C4/CA125 molecular subtype tumors. The observation that C1 tumors had higher stromal content is known, but the high concordance of tumor and stroma clusters with published HGSOC molecular subtypes suggested that genes which are differentially expressed between tumor and stroma might be sufficient to discriminate the AOCS molecular subtypes.

These molecular subtypes were defined using gene expression analysis, so we next attempted to quantify the relative contribution of stroma and tumor gene expression to each molecular subtype. We performed global differential gene expression analysis between each pair of AOCS tumor molecular subtypes (C1, C2, C4, and C5) (limma (26), 2-fold change and p<0.001 after FDR correction, Table S3). The number of genes that discriminated between each pair of subtypes varied in size from 343 (C1 *vs.* C2) to 1,267 (C1 *vs.* C5) (Figure 3A). We tested the overlap between these gene lists and the AOCS stromal and tumor gene sets. We observed that any contrast involving C1 had strong overlap with the stromal gene set (Fisher’s test p<10^−301^), as did the contrast of C2 *vs.* C4 (Fisher’s test p<10^−264^). Specifically, 53% and 41% of genes significantly different in contrasts of C1 *vs.* C2 or C1 *vs.* C4, respectively, were up-regulated in stromal cells (Figure 3A). This suggests that at least C1 is primarily driven by different amounts of stromal gene expression. In contrast, only 9% of genes differentially expressed between C4 and C5 were stromal genes, consistent with the subtypes’ reported low amount of desmoplasia (1). The tumor gene set and “other” genes dominated C5 gene differential expression, consistent with C5’s distinct molecular properties (2,35).

**Figure 3:**
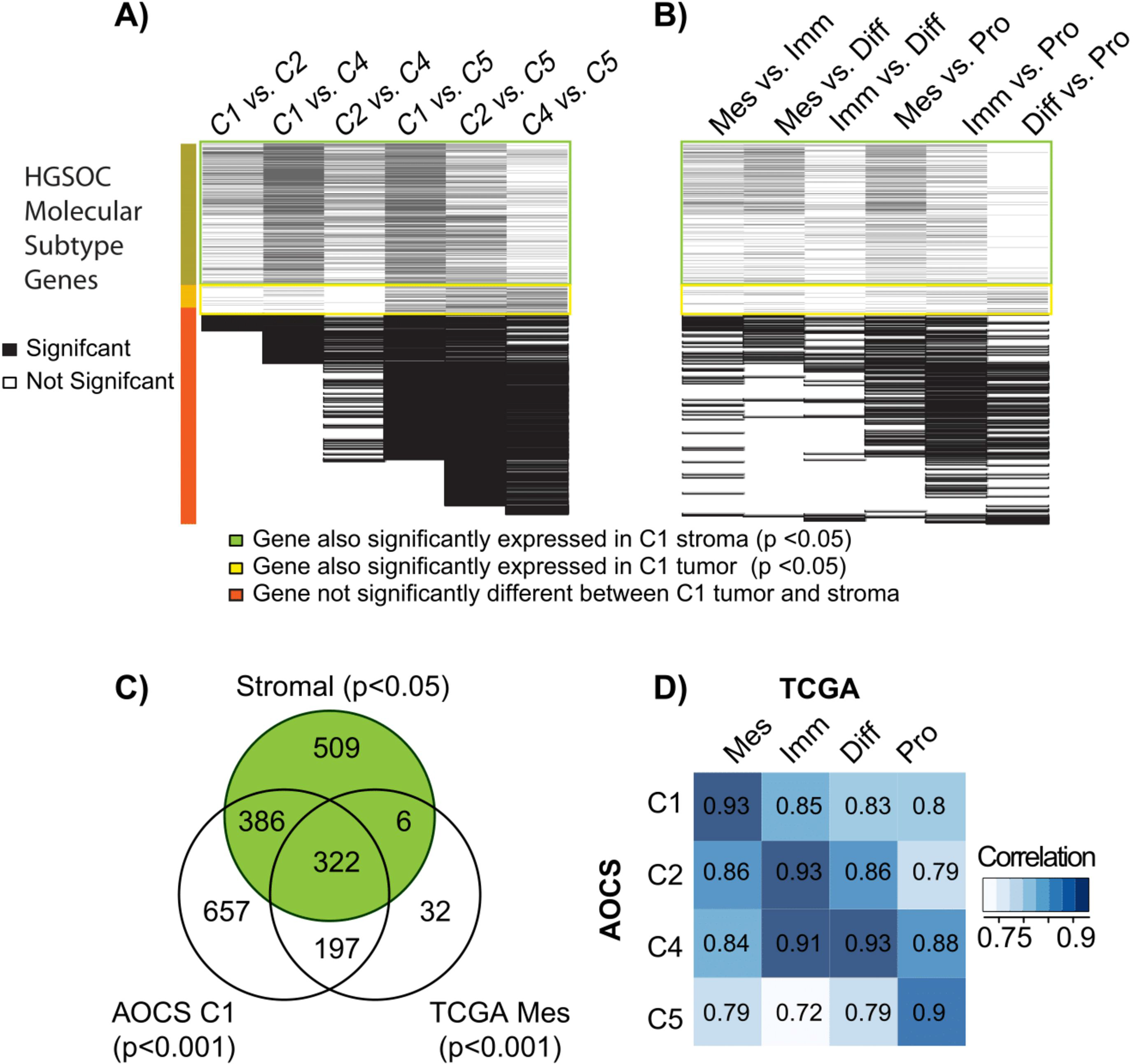
Overlap between tumor and stromal gene signatures and genes differentially expressed between the AOCS and TCGA molecular subtypes. Black and white lines indicate genes that were or were not strongly differentially expressed (limma, moderated *t*-statistics, p< 0.001 after FDR correction), respectively, between each pair of A) AOCS and B) TCGA molecular subtypes. The color bar indicates whether these genes were also differentially expressed in laser capture microdissected C1 tumor (yellow) and stroma (green) at p<0.05, or not significantly differentially expressed between C1 tumor and stroma (orange). Stromal and tumor genes with cutoff p<0.05 rather than 0.01 were used in order to explore the extent of stromal gene enrichment in these subtypes. C) There was a large overlap between genes expressed in C1 microdissected stroma and genes that were differentially expressed either between C1 vs. other AOCS subtypes, or between the TCGA mesenchymal subtype vs. other TCGA subtypes. D) Correlation between average expression of AOCS and TCGA molecular subtypes, which was calculated using the intersect of all differentially expressed genes between any pair of AOCS or TCGA subtypes. Mes, Imm, Diff and Pro are the TCGA molecular subtypes mesenchymal, immunoreactive, differentiated, and proliferative, respectively.

Based on these results, we also investigated the influence of stromal gene expression in TCGA HGSOC molecular subtypes. The TCGA mesenchymal, immunoreactive, differentiated, and proliferative subtypes have highest global gene expression correlation with the AOCS subtypes C1, C2, C4, and C5, respectively (Figure 3D), and the mesenchymal subtype has significantly higher percent stromal content than other subtypes (*t*-test p=1.35 × 10^−7^). Results from analysis of the TCGA study confirmed findings in the AOCS subtype analysis. Genes differentially expressed (p<0.001, FDR correction) in the TCGA mesenchymal subtype, similar to C1, were mostly in the stromal gene set (59%), and overlapped strongly (35%) with genes differentially expressed in C1 (Figure 3C). Similar to the AOCS C4 and C5 subtypes, few genes in the stromal gene set were differentially expressed between TCGA’s differentiated and proliferative molecular subtypes (<11%). Lastly, genes that were differentially expressed in the proliferative subtype, similar to C5, were dominated by tumor-specific and “other” genes.

### Molecular subtype classifications change with the proportion of stromal gene expression

We next investigated if the gene signature classifiers for these molecular subtypes were stable when the proportion of stromal cells changed. We trained a Single Sample Predictor (SSP) classifier using the gene lists reported to predict each of the AOCS C1, C2, C4, and C5 molecular subtypes (1), which predicted the training high-grade AOCS molecular subtypes with an overall self-validation accuracy of 95% (Figure S2A). The gene lists’ sensitivity and specificity for the molecular subtypes range from 0.93-0.97 and 0.96-1.0, respectively.

The AOCS molecular subtype classification was sensitive to percent stromal content. When applied to epithelial carcinoma cells microdissected from HGSOC tumors in an independent cohort (MGH dataset n=38), most microdissected tumors (20/38) were assigned to subtype C4 (Figure S2B). Matched microdissected stroma was all classified as C1 (n=23) or unclassified (n=15, Figure S2C). However, computational mixing of only 10% stromal gene expression with paired microdissected tumor caused 6/38 tumors to be reclassified (Figure S3), including C4 tumors that were reclassified as C1 (n=2) or C2 (n=2). Molecular subtype C5 tumors remained stable, reflecting the lack of stromal gene expression determining this subtype. Five additional tumors were predicted to have a different molecular subtype when the stromal content was increased to 20%, and at 30% stroma, over one third (15/38) of tumors’ subtypes were reassigned.

We observed similar results regarding the robustness of TCGA molecular subtypes with increasing proportions of stroma. We built an SSP classifier of TCGA molecular subtypes using their recently described 100-gene signature (7), which performed with an overall self-validation accuracy of 89%, with each subtypes’ sensitivity and specificity ranging from 0.84-0.94 and 0.95-0.98, respectively. Again, when applied to classify microdissected tumor in the MGH dataset, most tumors were assigned either to the differentiated subtype (23/38), which is similar to the AOCS C4 subtype, or the proliferative subtype (8/38), which is similar to C5 (Figure S2D). Microdissected stroma was classified as mesenchymal (20/38) or was unclassified (10/38, Figure S2E). With 10%, 20%, and 30% stroma, 0/38, 5/38, and 8/38 tumors, respectively, were classified to a different molecular subtype (Figure S3).

### The presence of stroma is weakly prognostic in HGSOC

We next explored whether percent stroma itself is prognostic. We first confirmed that the AOCS C1/stromal subtype (which has high percent stromal content) was associated with overall survival across all tumors (p=0.0083) and nearly significantly associated within high stage tumors (p=0.0538). In the TCGA data, pathologist scores of percent stromal content had a weak but significant (p=0.0337) association with overall survival in univariate Cox proportional hazards regression analyses of high stage (III or IV) tumors, but not across all tumors (p=0.0979). In other studies, no estimate of percent stroma was available, but we assessed the prognostic power of a tumor-stroma gene signature in an aggregated cohort of 2,527 HGSOC patients from 16 studies (modeled as fixed effects), which includes the TCGA and AOCS datasets (20). The 688-gene AOCS tumor-stroma signature was weakly prognostic with a hazard ratio (HR) of 1.17 (95% confidence interval 1.11-1.23, Figure 4A).

**Figure 4:**
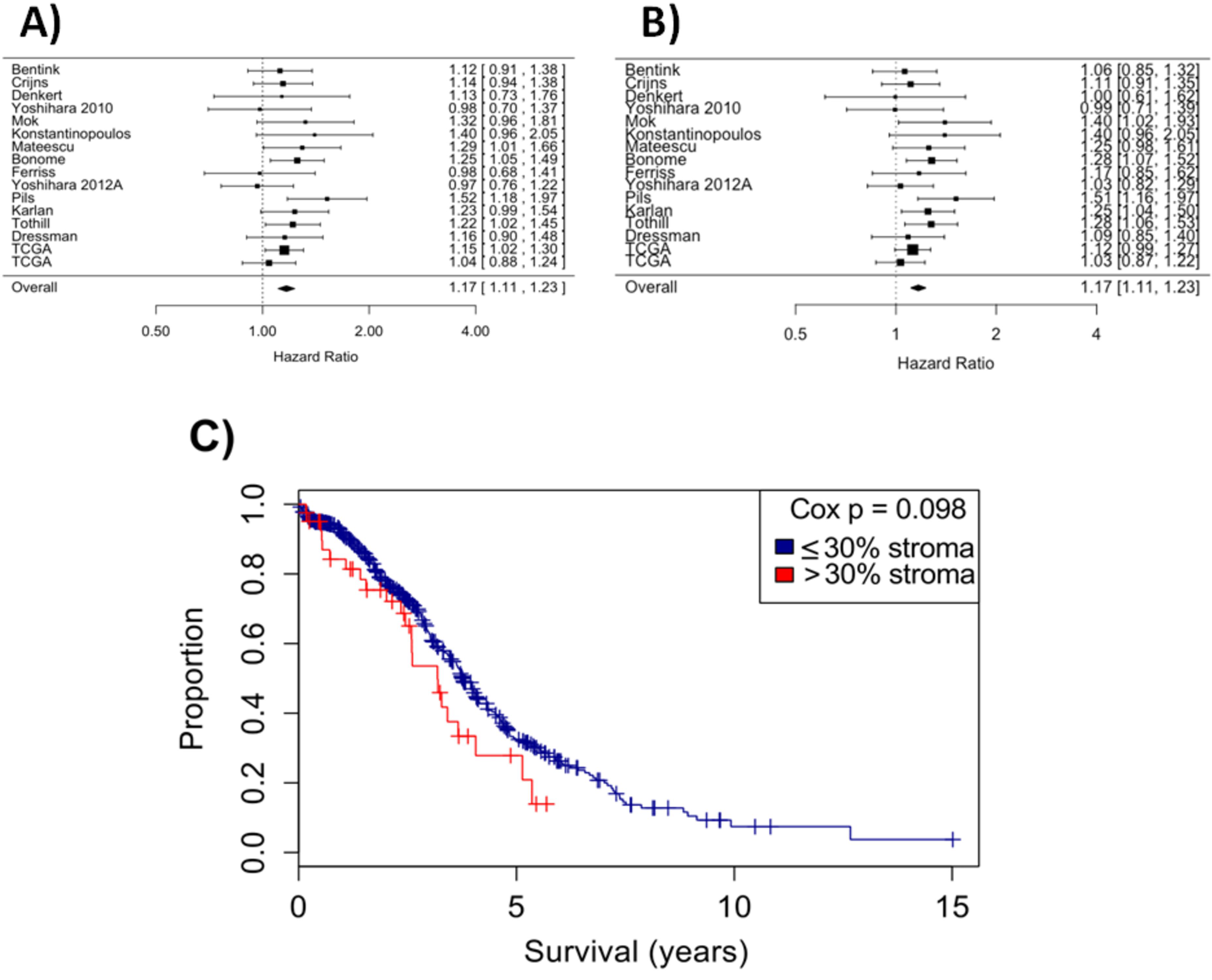
Stromal content is weakly prognostic of overall survival. Estimated stromal proportion using A) the AOCS tumor and stromal gene signatures or B only the genes that were specific to tumor stroma, which showed nearly identical prognostic power in sixteen ovarian cancer studies. C) Pathologist assessment of percent stroma in TCGA tumors was also weakly prognostic of overall survival.

It’s possible that the stromal genes are prognostic because of their expression within carcinoma cells, rather than because of their representation of stromal content, and thus we also examined genes that are only expressed in stroma. Using expression profiles of paired microdissected epithelial tumor (MGH dataset n=38), adjacent tumor stroma (n=38), and unrelated normal ovarian stroma (n=10), we applied a barcode approach (29) to convert gene expression to binary expressed or not expressed calls. Gene signatures of 28, 11 and 38 genes had greatest sensitivity and specificity (area under the receiver operator curve greater than 0.9 (30) Figure S4) for tumor, stroma adjacent to tumor, or normal ovary stroma (Table S2). The 11 genes specific to stroma had similar prognostic power (HR 1.17, CI 1.11-1.23, Figure 4B) but the 28-gene signature that was specific to tumor was not prognostic (HR 0.96, CI 0.96-1.01, data not shown) and performed comparably to a 20 gene-randomly selected gene signature (HR 0.96, CI 0.96- 1.02, data not shown) in the meta-analysis of 2,527 tumors, which supports the involvement of stroma in patient outcome.

### Several prognostic ovarian cancer signatures are correlated with percent stroma

Given prognostic ability of stroma in HGSOC, we hypothesized that published prognostic ovarian gene signatures might be a proxy for the presence of stromal tissue. Published gene signatures of HGSOC (n=61) were downloaded from the database GeneSigDB (32) or curated from the literature (Table S4). Gene signatures varied in size from four to 1,903 genes, with a median of 47 genes. Over a third of signatures (n=24/61) were enriched in the AOCS stromal genes, and 11/61 were enriched in MGH stroma genes (p<0.05 after FDR correction). Eight signatures (AOCS signatures; Biade et al. benign tumor signature (36); Bentink et al. angiogenic signatures (11); and prognostic signatures from Bignotti et al. (37), Bonome et al. (38), and Spentzos et al. (39)) were strongly enriched in stromal genes (p<0.001) in both analyses (Table S4) and were examined in greater detail (Table 1). Other gene signatures did not significantly overlap with the stromal gene set, including the TCGA study’s prognostic signature (Table S4) and the prognostic BRCAness signature.

**Table 1:**
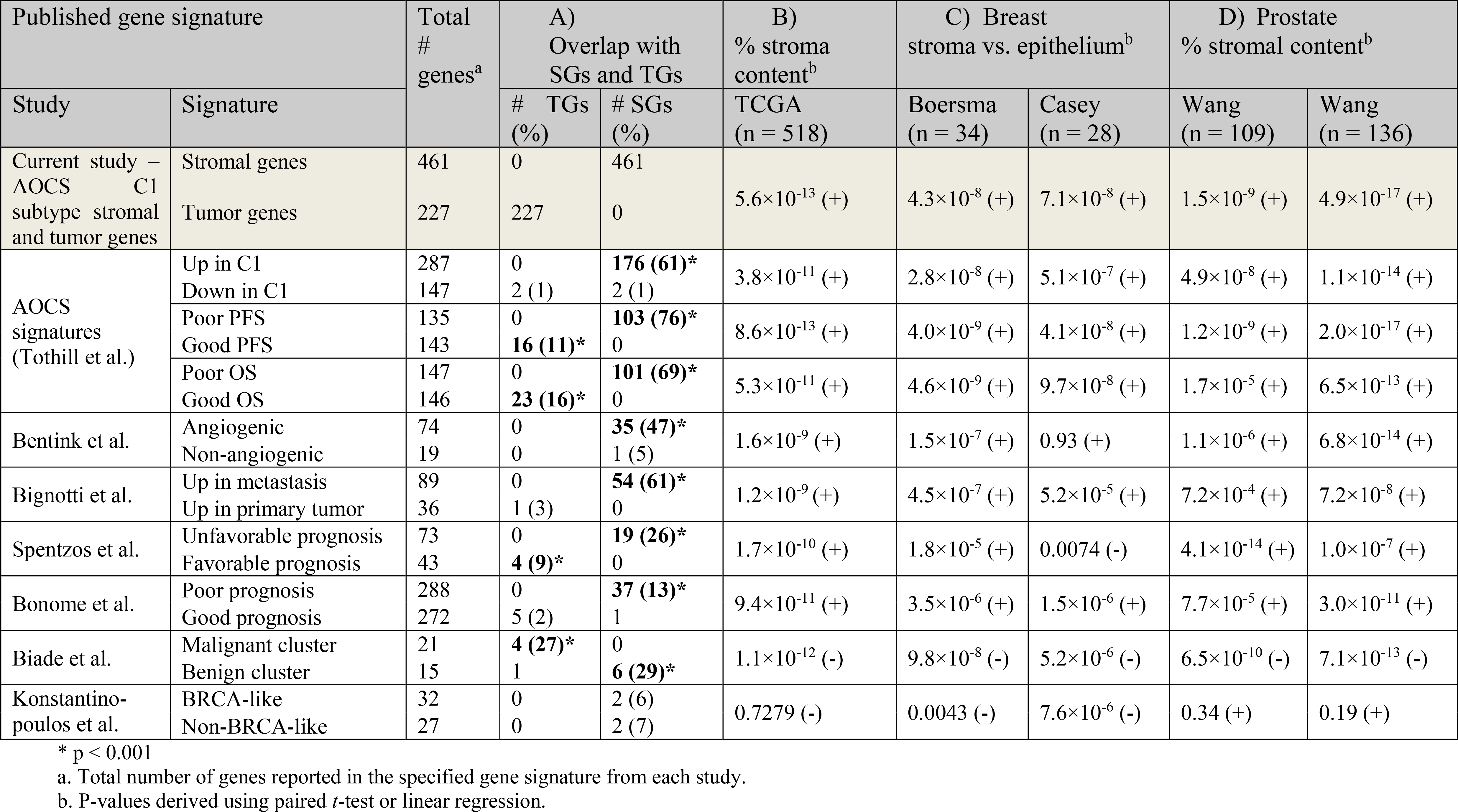
Comparisons between the ovarian cancer gene signatures and stromal genes, tumor genes, or percent stromal content. A) We calculated the overlap between several ovarian cancer gene signatures and the stromal gene (SG) set and tumor gene (TG) set derived from the AOCS dataset. P-values for these columns were calculated with Fisher’s exact test. B-D) We also listed p-values for a signature’s association with either percent stromal content or microdissected stroma in B) the TCGA ovarian cancer dataset, C) two breast cancer datasets, and D) two prostate cancer datasets. In B-D), p-values were calculated using linear regression or /-tests. Of the two components for each signature, the component listed first had the +1 weight for the signature’s score (e.g. genes “Up in C1” have a +1 coefficient, and genes “Down in C1” have a −1 coefficient). Directions of association (+/−) with stroma are listed in parentheses next to p-values. No multiple testing correction was used.

In published gene signature with significant overlap with tumor and stromal gene signatures, the poor prognosis genes overlapped with the stromal gene set in 7/8 gene signatures (p<0.001), and the good prognosis components were enriched in tumor genes in several of these signatures (Table 1A). The eight published gene signatures that were enriched in stromal gene expression were also positively associated with pathological scores of percent stromal content in TCGA tumors (linear regression, p<10^−8^, Table 1B).

Of note the Biade *et al.* signature (36) was an exception because stromal gene expression was enriched in expression profiles of benign tissue, although this is consistent with their reports of increased stromal cell proportions in their benign tumor samples (36). The Konstantinopoulos et al. BRCAness signature (40) was not significantly enriched for either stromal or tumor genes it and was not associated with percent stromal content in TCGA samples, so it represented a negative control.

### The stromal signature is not specific to HGSOC

Many epithelial tumors have complex interactions with other cells in the tumor microenvironment. Therefore, we asked whether these stroma-associated ovarian cancer gene expression profiles are specific to HGSOC, or might more broadly characterize epithelial cancers. We found that ovarian cancer stromal genes significantly predicted tumor *vs.* stroma or were associated with the percentage of stroma in other epithelial cancers. Each of the eight ovarian cancer signatures enriched in stromal genes was associated with breast (Boersma et al. (21), Casey et al. (22)) and prostate (Wang et al. (9)) stroma in at least three of four datasets tested (linear regression or paired *t*-test p<10^−4,^ breast Table 1C, prostate cancer Table 1D).

### Clinical variables that explain prognostic power of stroma-associated genes

Lastly, we explored what variables may explain the prognostic power of these published gene signatures. Most gene signatures that were enriched in stromal genes were no longer associated with poor overall survival (OS) when adjusted for pathologist’s estimates of percent stromal content in TCGA tumors in Cox proportional hazard regression analysis (Table 2A). By contrast, the BRCAness signature remained prognostic of OS in multivariate models that were adjusted for both stromal content and stage (p<0.05, Table 2A). We further observed that the AOCS prognostic gene signatures remained associated with OS and progression free survival (PFS) when adjusted for stromal content (p<0.01 and p<0.05, respectively, Table 2A), but were no longer prognostic after adjusting for both stromal content and stage.

**Table 2:** Association between each ovarian cancer gene signature and overall survival. A) The table lists each signature’s score’s p-value for association with overall survival (OS) in the TCGA dataset, with p < 0.05 in bold. These associations were also examined in multivariate analyses after adjusting for percent stromal content and stage. B) The signatures were examined in the smaller MGH dataset, applying signatures to either the microdissected epithelium or stroma.

| Published gene signature |  | A) HGSOc tumors (TCGA, n=518) |  |  |  | B) MGH Data (n=38) |  |
| --- | --- | --- | --- | --- | --- | --- | --- |
| Study | Signature or % stromal content | OS | OS, adjust stage | OS, adjust % stroma | OS, adjust % stroma + stage | OS, microdissected epithelium | OS, microdissected stroma |
| AOCS C1 stroma vs. tumor<sup>a</sup> | Stromal genes vs. tumor genes | 0.0870 | 0.137 | 0.2730 | 0.484 | 0.2440 | 0.1800 |
| MGH stroma vs. tumor<sup>a</sup> | Activated and normal stroma vs. tumor | **0.0336** | 0.0544 | 0.148 | 0.289 | 0.555 | 0.156 |
| MGH tissue-specific genes<sup>b</sup> | Tumor | 0.778 | 0.640 | 0.996 | 0.906 | 0.924 | 0.718 |
|  | Tumor Stroma | **0.0430** | 0.0732 | 0.177 | 0.337 | 0.540 | 0.164 |
|  | Normal or Tumor Stroma | **0.0395** | 0.0707 | 0.140 | 0.294 | 0.598 | 0.175 |
| TCGA stromal content | Pathologist assessment of % stroma in TCGA samples | 0.0979 | **0.0499** | N/A | N/A | N/A | N/A |
| AOCS (Tothill et al.) | C1 | **0.0175** | 0.0658 | 0.0755 | 0.276 | 0.343 | 0.316 |
|  | PFS | **0.0045** | **0.0195** | **0.0346** | 0.148 | 0.242 | 0.377 |
|  | OS | **0.0010** | **0.0068** | **0.0076** | 0.0541 | 0.356 | 0.105 |
| Bentink et al. | Angiogenic vs. non-angiogenic | **0.0313** | 0.115 | 0.109 | 0.374 | 0.552 | 0.324 |
| Bignotti et al. | Metastasis vs. primary | **0.0165** | 0.0921 | 0.0613 | 0.311 | 0.383 | 0.229 |
| Spentzos et al. | Prognosis | **0.0118** | **0.0330** | 0.0507 | 0.163 | 0.608 | 0.145 |
| Bonome et al. | Prognosis | **0.0426** | 0.0571 | 0.128 | 0.208 | 0.366 | **0.0352** |
| Biade et al. | Malignant vs. benign | 0.310 | 0.266 | 0.587 | 0.584 | 0.642 | 0.101 |
| Konstantinopoulos et al. | BRCA-like | **0.0167** | **0.0281** | **0.0272** | **0.0386** | 0.696 | 0.387 |

<sup>a</sup> Table S1, <sup>b</sup> Table S2

We hypothesized that the difference in stroma’s prognostic power between these two datasets was related to heterogeneity in sampling location. In the AOCS C1 subtype, 61% of tumor samples were from extra-ovarian sites (most commonly peritoneum, as defined by the AOCS study), which often reflects a higher stage and thus worse prognosis, compared to 19% extra-ovarian sampling in other AOCS subtypes (1). In the TCGA dataset, every tumor except for one was reported to be sampled at the ovary/pelvis (2). After adjusting for sampling location, C1’s association with disease-free survival (adjusted p=0.03) and overall survival (adjusted p=0.06) were substantially attenuated. Thus, C1’s prognostic power is partially associated with sampling location.

## Discussion

We show that the proportion of stroma admixture in bulk tumor is a source of differential gene expression that impacts molecular subtyping and survival analysis in HGSOC. Furthermore, a subset of published prognostic gene signatures of ovarian cancer are enriched in stromal genes and are strongly associated with the proportion of stroma, and their association with overall survival is no longer significant after adjusting for percent stroma in bulk HGSOC tumors.

The proportion of stromal cells determined by pathology inspection was a weak predictor of poor overall survival in HGSOC patients. Pathologist assessments of stromal proportion are available in few gene expression studies, so we estimated stroma proportion using gene signature models and found that stroma proportion was weakly prognostic in a meta-analysis of 2,527 tumors from independent gene expression studies (HR 1.17) (3,4). A recent analysis of TCGA tumor purity found an association between ovarian cancer tumor purity and lymphatic invasion but did not report an association with survival (41). That study examined a consensus measurement of purity estimations (CPE), which was the median normalized value of four methods, two of which specifically estimated immune infiltrate. That study did not focus on tumor-associated stroma, and we found that CPE estimates (41) were negatively correlated with pathologist estimates of stromal proportion in HGSOC TCGA tumors (Pearson Correlation - 0.84).

The prognostic performance of stroma proportion could be partly explained by an association with stage or sampling location. In the AOCS, proportion of stroma was associated with sampling location; 40% of C1 tumors (in contrast to 9% in other subtypes) were reported to contain ≥50% stromal content, and this molecular subtype also had higher numbers of tumors that were obtained from the peritoneum or other metastatic sites. This association with sampling location may reflect site-specific or intra-tumor variability in the proportion of stroma. These sources of variability are an important considerations in molecular subtype and prognostic biomarker discovery, given that a high proportion of stroma was not always associated with more advanced tumors, such as in the benign cluster reported by Biade (36).

Stromal gene expression substantially influences “observed” gene expression and impacts tumor molecular subtype discovery. Small increases in stroma gene expression caused subtype misclassification in our computational model that mixed stroma and tumor gene expression profiles. Although adjacent stromal content can affect an individual epithelial cell’s gene expression pattern (42), this is distinct from changes in observed gene expression caused by the admixture of stromal cells and epithelial carcinoma cells in bulk tumors. We show that stromal genes differentiated the AOCS C1 molecular subtype or the TCGA mesenchymal subtype from other HGSOC molecular subtypes. In an miRNA analysis, the mesenchymal TCGA molecular subtype overlapped with the proliferative subtype, and the mesenchymal TCGA molecular subtype lacks a reproducible genetic phenotype (gene mutation or copy number alternations) (2). The biological basis for the increased stroma proportion in these subtypes is unknown, and tissue sampling heterogeneity needs to be considered since the C1 samples were enriched in tumors sampled from the peritoneum. In contrast, the AOCS C5 subtype and the similar TCGA proliferative subtype are reported to have genetic alterations, including copy number variation, in several genes (2,35), and AOCS C5 subtype gene expression patterns were not well explained by stromal genes. The impact of tissue sampling heterogeneity in molecular subtype discovery of gene expression profiles of bulk tissue were recognized as early as 1999 when Golub et al. (43) remarked: “studies will require careful experimental design to avoid potential experimental artifacts ‒ especially in the case of solid tumors. Biopsy specimens, for example, might have gross differences in the proportion of stromal cells. Furthermore, accumulating evidence suggests that the composition and gene expression of stroma may vary depending upon the specific anatomic location of the specimens. Blind application of class discovery could result in identifying classes reflecting the proportion of stromal contamination in the samples, rather than underlying tumor biology. Such ‘classes’ would be real and reproducible, but would not be of biological or clinical interest.”

The influence of non-cancer cell gene expression on subtype classification applied to bulk tumor profiles is potentially a source of systematic error in several solid tumors. The accuracy of signature classifiers that predict breast tumor molecular subtype (PAM50) or risk of recurrence (Oncotype DX) decrease considerably with decreasing proportion of tumor cells and increasing proportion of normal cells (34). Molecular subtype definitions developed from gene expression profiles of bulk ovarian, breast, and other carcinomas may be redefined when molecular subtypes of both tumor and stroma are considered individually. Similar to studies of immune molecular subtypes across a wide array of different cancers (28,44), there is a need to define the landscape of stromal heterogeneity in carcinomas. Thus far, at least two HGSOC stroma subtypes have been proposed (13). Deconvolution of cell mixtures in gene expression studies has also been an active area of research (45), and towards this end, we identified tumor- and stroma-specific genes of HGSOC epithelial cancer cells and stromal cells (Table S2) that may be applied to computational predictions of cell proportions and further studies of tumor/stroma interactions.

These tissue-specific genes also highlight the importance of studying both the cancer cell and stromal cell biology. The gene CEP55 had the highest specificity and sensitivity of the 28 tumor-specific genes, and it codes for a protein that interacts with BRCA2 and plays a critical role in regulating the final step of cytokinesis (46). Two of the eleven stroma-specific genes (IFFO1, GLIPR1) were strong predictors both of pathologist estimates of stroma proportion in TCGA tumors (Pearson correlation 0.7 and 0.67, respectively, p<0.0001) and of a gene signature that predicts macrophage/monocyte activity (Figure S4, colony stimulating factor-1, CSF1 signature) (28,47). Another three stroma-specific genes were strongly correlated with TGCA immune landscape scores for TGF-β response (Figure S4) in TCGA tumors (28). These three genes included KCNE4; Thrombospondin 1 (THBS1), which is associated tumor growth and metastasis in gastric carcinoma (48); and biglycan (BGN), which has been shown form a complex with either TGF-β1 or TGF-β1 type I receptor to intensify the phosphorylation of Smad2/3 in cultured endothelial cells (49). These two subgroups in the 11 stroma-specific genes are potentially related to the recently described TGF-β-dependent or TGF-β-independent Smad signaling (13) in stroma of HGSOC and support the need for further study of stromal subtypes.

Our results strongly support single cell analysis or microdissection of tumor samples in gene expression studies so that the features of the epithelial tumor cells, stromal, and immune components can be each analyzed. Retrospective analysis of bulk gene expression patterns in light of emerging single cell sequencing data are needed to refine molecular subtypes in ovarian cancer and other carcinomas and enable a more comprehensive search for molecular targets amenable to therapeutic intervention.

## Supporting information

## Supplementary Tables

**Table S1: Datasets from the curatedOvarianData R package.** In this package, the stage column provides early, late, or unknown, and the histology column provides serous, clear cell, endometrioid, mucinous, other, or unknown.

**Table S2: Genes differentially expressed between stroma and tumor.** These include genes up in C1 tumor compared to stroma (tumor gene set) and genes up in C1 stroma compared to tumor (stromal gene set) at A) p<0.01 and B) p<0.05, as well as genes C) differentially expressed between MGH tumor and stroma. D) Genes specific to tumor, tumor-stroma, or tumor or normal stroma, which were derived using the MGH dataset. For this MGH analysis, gene expression values were converted to binary on/off using the barcode method (29), and to identify genes with greatest sensitivity and specificity for tissue types, only genes with receiver operating curve (ROC) > 0.9 were considered.

**Table S3: Genes differentially expressed between ovarian cancer subtypes.** These are lists of genes that are differentially expressed between each of four high-grade ovarian cancer subtypes in both the AOCS dataset and the TCGA dataset at p<0.001 using modified t-test and FDR correction (25) in Bioconductor package limma (26).

**Table S4: Complete list of all ovarian cancer gene signatures that were analyzed, as well as their enrichment in the tumor-stroma gene signatures.** Fifty signatures were from GeneSigDB, and the other signatures, except for the Konstantinopoulos et al. “BRCAness” signature and TCGA study’s prognostic gene signature, were chosen for analysis because of their reported association with the C1 subtype. P-values were corrected for multiple testing using the Benjamini-Hochberg method (25).

## Supplementary Figures

**Figure S1:**
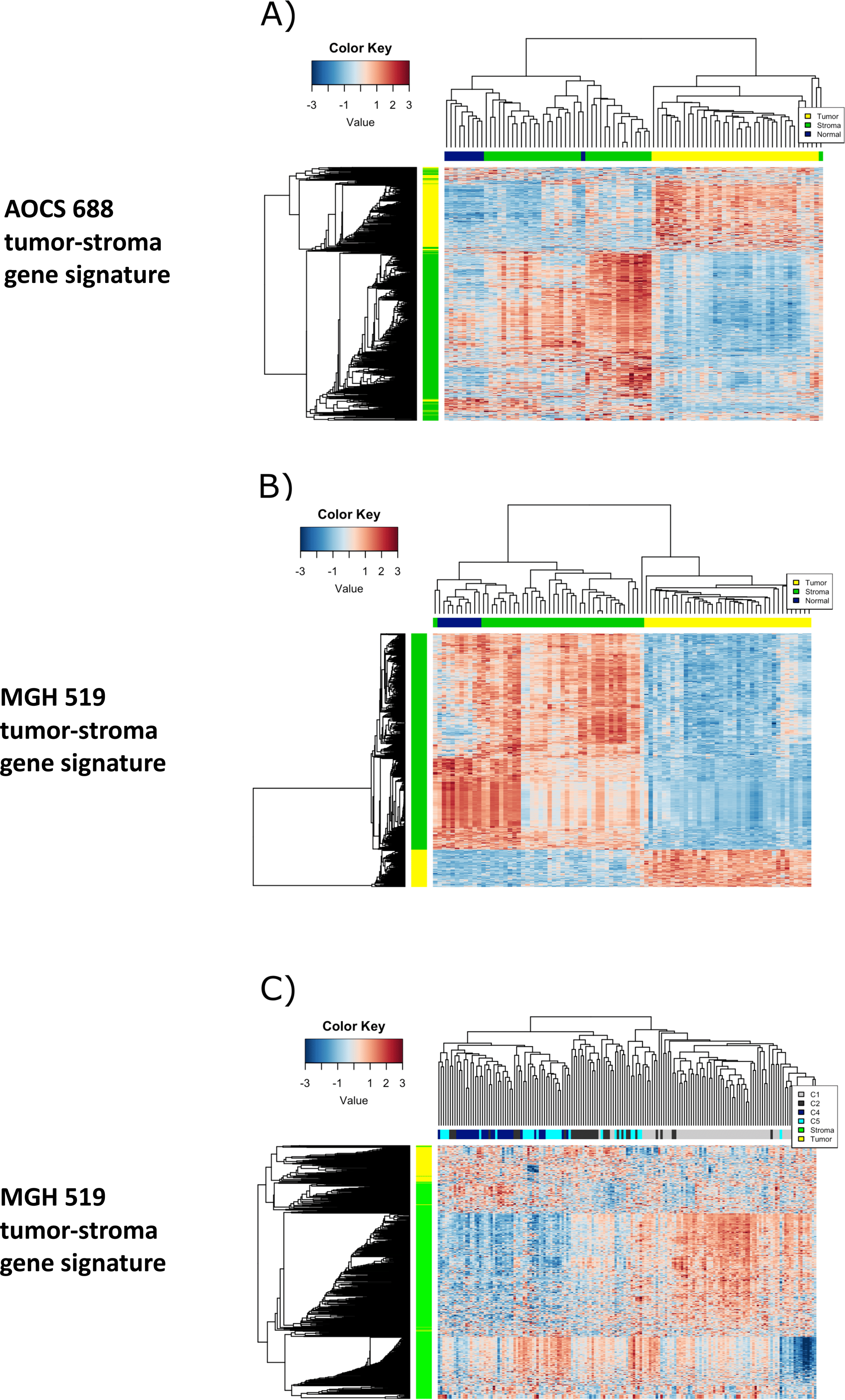
The AOCS and MGH signatures discriminate tumor and stroma tissue and separate serous ovarian cancer molecular subtypes. Heatmap of the gene expression of A) the AOCS 688 tumor-stroma gene signature (developed on the AOCS C1 microdissected tissue) in an independent MGH dataset, which shows that it partitioned all but one of the 38 pairs microdissected tumor and stroma MGH samples. Similarly, the gene expression of the 519 genes MGH tumor-stroma gene signature distinguished the microdissected tumor and stroma in B) the MGH tumors (training dataset) and C) the AOCS tumors where 2 large clusters were observed, which distinguished molecular subtypes C1 from the other AOCS molecular subtypes. The second cluster, could be split into two clusters which were enriched in C2 and C4/C5 tumors respectively. All of these analyses were done using unsupervised hierarchical cluster analysis. Colors of the heatmap were scaled using z-score values across rows.

**Figure S2:**
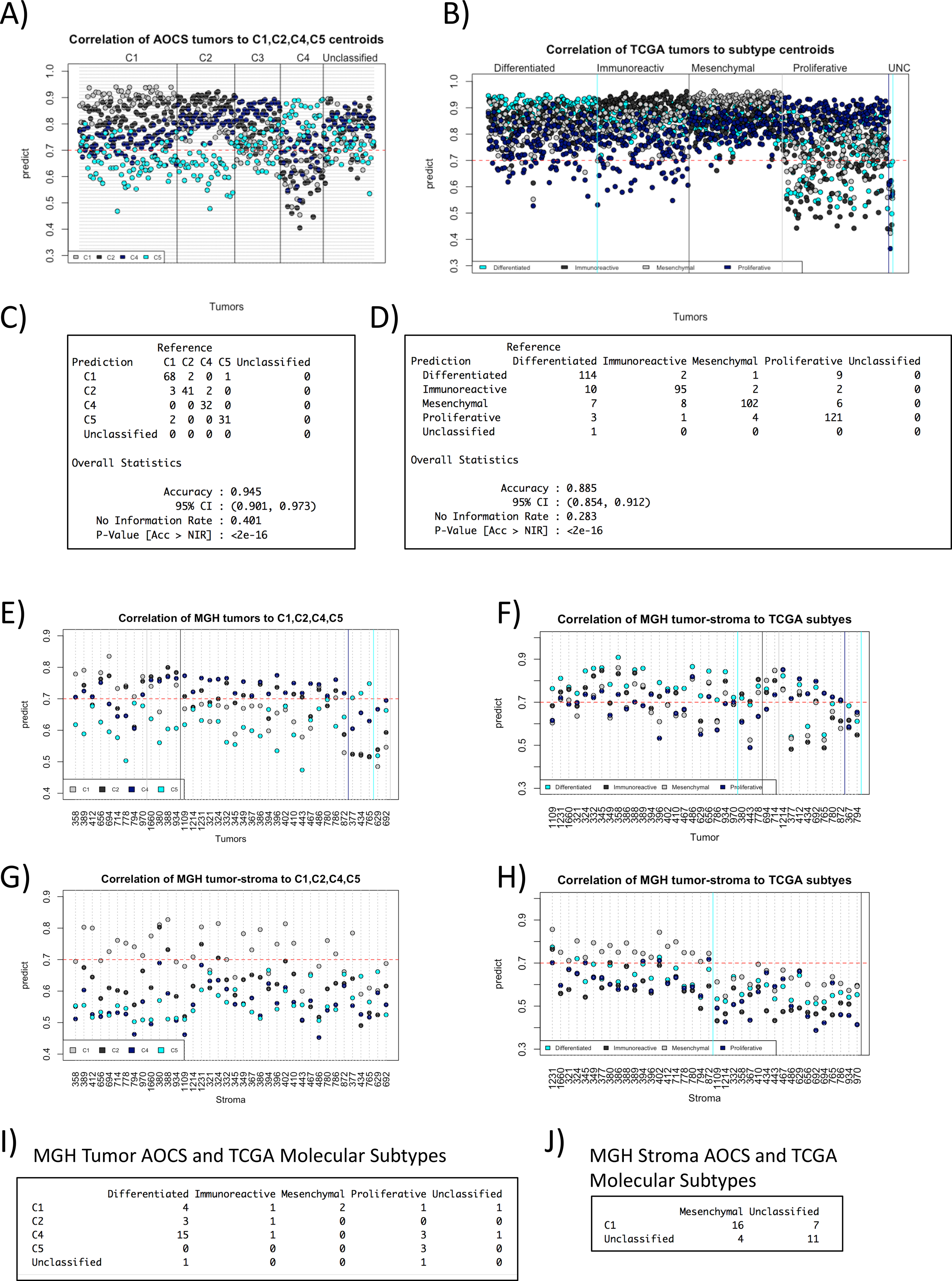
Training ovarian cancer molecular subtype classifiers. Single sample predictors (SSP) of the AOCS (A) and the TCGA (B) molecular subtype classifiers were developed, and tumors were assigned a molecular subtype based on the highest Pearson correlation coefficients to each subtype’s centroid. Figures A) and C) show the correlation of AOCS subtypes to AOCS subtype centroids, and B) and D) the TCGA tumors to TCGA subtype centroids. For example, in B) we see 68 C1 tumors had highest correlation to C1 clusters and were assigned to C1. The performance of the AOCS and TCGA SSP classifiers on their training data is provided in B) and D) respectively. The molecular subtypes are color-coded: C1 (light grey), C2 (dark grey), C4 (navy), and C5 (cyan). Microdissected tumor (E,F) and stroma (G,H) were assigned to AOCS (E,G) or TCGA (F,H) molecular subtypes. A large number microdissected MGH tumors (n=15/38) were classified as ACOS C4 or the TCGA differentiated molecular subtype, whereas most microdissected MGH stroma samples were classified as C1/Mesenchymal (16/38) or unclassified (11/38).

**Figure S3:**
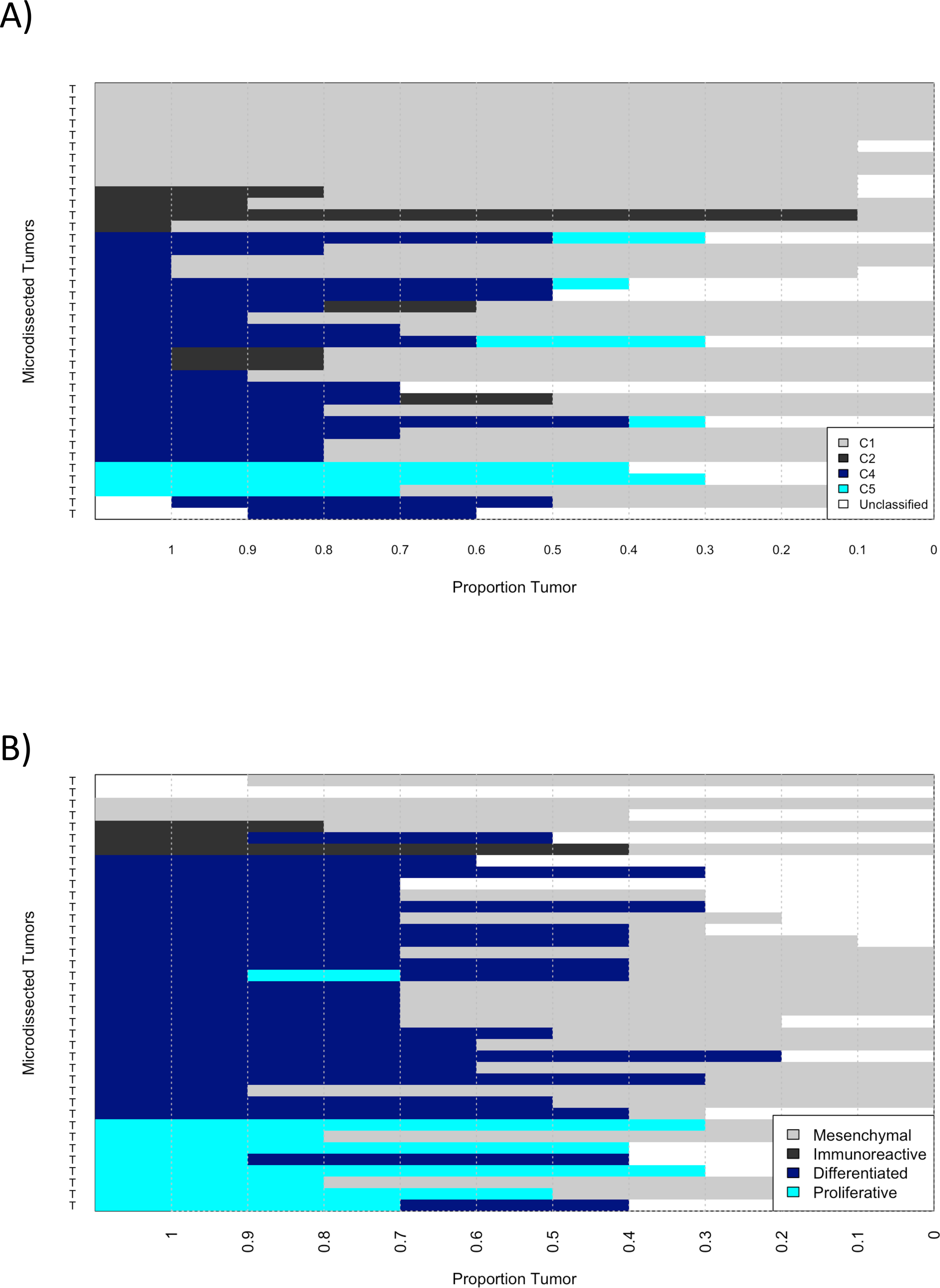
Reclassification of samples with increasing stroma proportions. MGH microdissected tissue was classified into molecular subtypes after computational mixing of paired microdissected tumor and stroma tissue, using 10% increments of stroma. MGH tumors are reclassified using paired MGH stroma samples and A) the AOCS subtype classification or B) the TCGA subtype classification.

**Figure S4:**
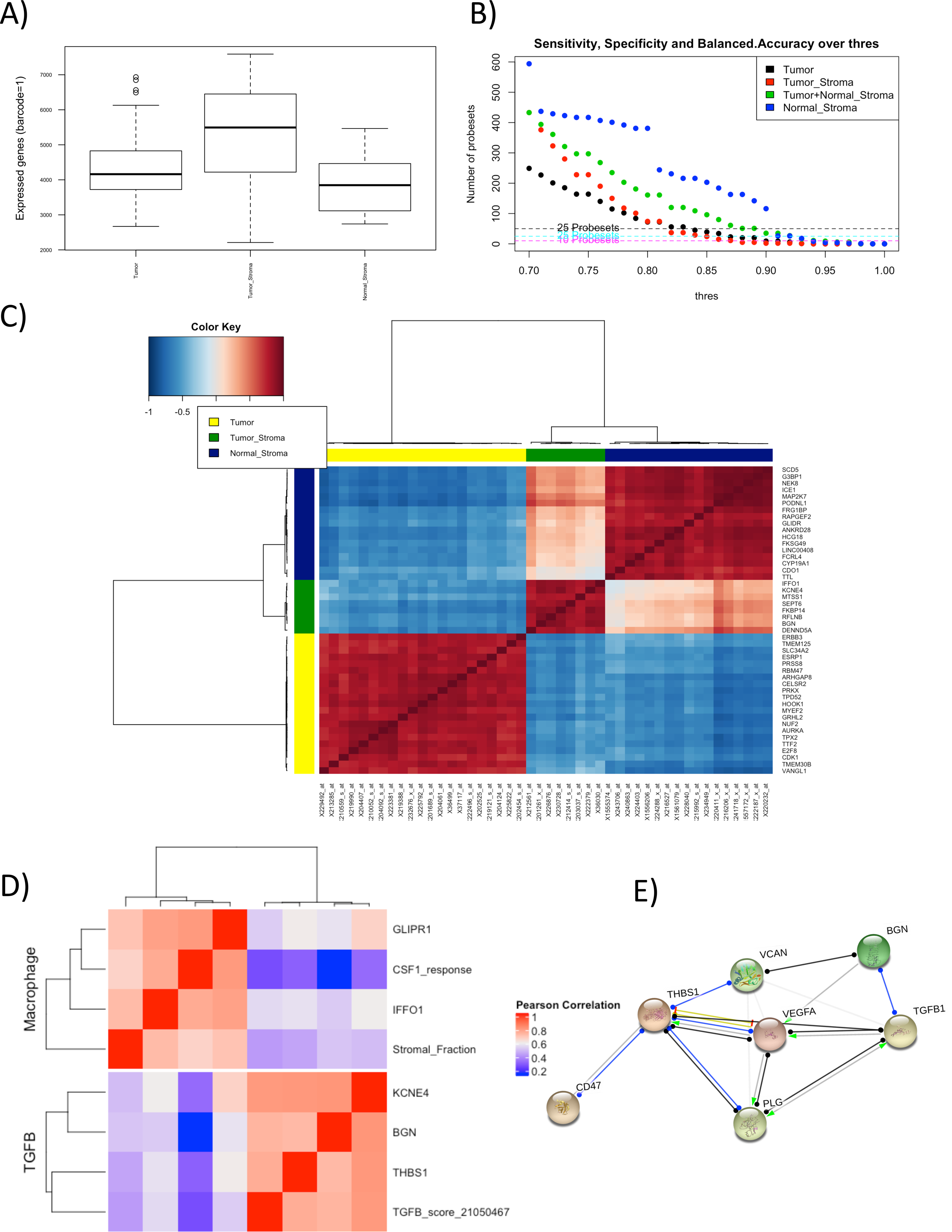
Discovery of tissue-specific gene signatures in microdissected tumor and stroma. The barcode method was used to discretize gene expression values into 0 and 1 calls. 34706 (63.48%) of Affymetrix probeIDs were not expressed in any sample and A) 19969 (36.52%) of Affymetrix probe IDs were expressed (barcode =1) in at least one tissue, with more probeIDs expressed in tumor-associated stroma (tumor-stroma), possibly reflecting the heterogeneity of cell types in tumor-stroma. We identified genes with greatest tissue sensitivity and specificity (area under the receiver (AUC) operator curve greater than 0.9) for tumors, tumor-stroma, or normal ovarian stroma. This resulted in 28, 15, 82, and 205 genes that had an AUC of at least 0.9 for tumor, tumor-stroma, tumor-stroma/normal stroma combined, or normal stroma, respectively. Probes (n=4) that were found in tumor-stroma and normal stroma were excluded. An AUC of 0.95 was used for normal stroma specifically, reducing it to 205 probe IDs. Figure C) shows a heatmap of the gene-gene correlations in gene expression profiles of MGH microdissected tissue for those genes with AUC >0.91. Figure D) shows the tumor-stroma genes with correlations >0.9 to Immune landscape gene sets scores of HGSOC tumors (28). This heatmap suggested subtypes, with genes related to TGF-β and others independent of TGF-β (macrophage/CSF1 subtype). Figure E) shows theoretical interactions between several tumor-stroma genes and genes extracted from the StringDB (https://string-db.org/) database (50).

